# Serine synthesis pathway upregulated by E-cadherin is essential for the proliferation and metastasis of breast cancers

**DOI:** 10.1101/2023.05.24.541452

**Authors:** Geonhui Lee, Claudia Wong, Anna Cho, Junior J. West, Ashleigh J. Crawford, Gabriella C. Russo, Bishwa Ranjan Si, Jungwoo Kim, Lauren Hoffner, Cholsoon Jang, Moonjung Jung, Robert D. Leone, Konstantinos Konstantopoulos, Andrew J. Ewald, Denis Wirtz, Sangmoo Jeong

## Abstract

The loss of E-cadherin (E-cad), an epithelial cell adhesion molecule, has been implicated in the epithelial-mesenchymal transition (EMT), promoting invasion and migration of cancer cells and, consequently, metastasis. However, recent studies have demonstrated that E-cad supports the survival and proliferation of metastatic cancer cells, suggesting that our understanding of E-cad in metastasis is far from comprehensive. Here, we report that E-cad upregulates the *de novo* serine synthesis pathway (SSP) in breast cancer cells. The SSP provides metabolic precursors for biosynthesis and resistance to oxidative stress, critically beneficial for E-cad-positive breast cancer cells to achieve faster tumor growth and more metastases. Inhibition of PHGDH, a rate-limiting enzyme in the SSP, significantly and specifically hampered the proliferation of E-cad-positive breast cancer cells and rendered them vulnerable to oxidative stress, inhibiting their metastatic potential. Our findings reveal that E-cad adhesion molecule significantly reprograms cellular metabolism, promoting tumor growth and metastasis of breast cancers.

## Introduction

E-cadherin (E-cad), encoded by the *CDH1* gene, is a key molecule for adherens junctions between epithelial cells. As it plays a critical role in tissue barrier formation and organ homeostasis [1–3], its dysregulations are directly associated with various diseases, including cancer. In particular, inactivating mutations of *CDH1*, consequently leading to loss of E-cad expression, were observed in most invasive lobular breast cancers [4]. Also, the epithelial-mesenchymal transition (EMT), an initiating step of metastasis, is often associated with loss of E-cad in multiple cancers [5, 6]. However, recent studies have demonstrated that E-cad could promote tumor progression and metastasis. For example, E-cad expression is strongly correlated with a worse prognosis in invasive ductal carcinomas, the most common type of breast cancers, and pancreatic cancers [7, 8], and it is frequently found in metastatic foci in patient samples [9]. Furthermore, E-cad upregulation accelerates tumor growth by activating signaling pathways [10] and elevates metastatic potential by reducing reactive oxygen species (ROS) levels [7]. To meet anabolic needs or manage oxidative stress, cancer cells must upregulate specific metabolic pathways. Still, our understanding of how E-cad confers metabolic advantages on cancer cells is limited.

Here, we report that the *de novo* serine synthesis pathway (SSP) plays an essential role for the proliferation and metastasis of E-cad-positive breast cancers. The SSP, a metabolic branch from the glycolysis pathway, contributes to lipid and nucleotide syntheses [11], redox homeostasis [12], and TCA anaplerosis [13]. Therefore, SSP upregulation is associated with aggressiveness and drug resistance of various cancers, including breast and liver cancers [14, 15]. We find that E-cad positively regulates the expression of phosphoglycerate dehydrogenase (PHGDH), the first and rate-limiting enzyme in the SSP, and targeting PHGDH abrogates the metabolic advantages conferred by E-cad in multiple breast cancer cells. Also, we show that E-cad-mediated PHGDH upregulation requires c-Myc, a major transcription factor that directly regulates multiple metabolic pathways. Our results reveal a novel molecular link between the adherens junction molecule E-cad and cell metabolism and illustrate that E-cad-mediated metabolic reprogramming is essential for tumor growth and metastasis.

## Materials and methods

### Antibodies

PHGDH (#13428), c-Myc (#5605S), PDH (#3205S), phospho-PDH (#31866), Smad2/3 (#8685S), HIF-1α (#14179S), HIF-2α (#7096S), ASCT2 (#8057) and E-cadherin (#3195S, #14472S) antibodies were obtained from Cell Signaling Technology for immunoblot or fluorescence imaging analysis. PSAT1 (#NBP1-32920, Novus Biologicals), PSPH (#PA522003, ThermoFisher), SIRT2 (#09-843, Millipore Sigma), MCT1 (#20139-1-AP, ThermoFisher), MCT4 (#22787-1-AP, Proteintech), and β-Actin (#ab8226, Abcam) antibodies were obtained for immunoblot analysis. HECD-1 (#13-1700, ThermoFisher) antibody was obtained for E-cad functional blocking.

### Proteomic data analysis

The proteomic data were quantified using log2 transformation and median normalization to the pooled reference channel (TMT-126) with an additional normalization step including the median of each sample. List of metabolic pathways and protein-coding genes were obtained from the KEGG database (https://www.kegg.jp/kegg/). Heatmap and volcano plot were generated with R Studio (https://www.r-project.org/) using pheatmap package [54] and Enhanced Volcano package [55]. Principal component analysis (PCA) and Gene Ontology (GO) analysis were obtained using the DESeq2 via DEBrowser using default parameters [56]. Gene expressions of the proteomic data were considered differentially expressed with FDR of < 0.05 and log2 fold change of ±0.58 (larger than 1.5-fold).

### Determination of metabolic pathway activity

To determine the *i*-th metabolic protein-coding gene expression of the replicated batch number in *j*-th metabolic protein-coding genes, we first calculated fold change:

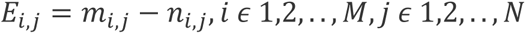

where *E*_*i,j*_ is the log2 transformed fold change of protein-coding genes of the *i*-th cell of E-cad^+^ or E-cad^-^ cell in the *j*-th metabolic protein-coding genes, *m*_*i,j*_ is the log2 transformed protein-coding gene expression of E-cad^+^ cell, *n*_*i,j*_ is the log2 transformed protein-coding gene expression of E-cad^-^ cell, M is the replicated batch number of proteomics data, and N is the number of metabolic protein-coding gene of the metabolic pathway. We then determined the metabolic activity using averages of each fold change of protein-coding genes:

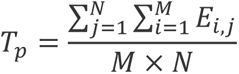

where *T_p_* is metabolic activity of *p*-th pathway which include metabolic gene categorized by the KEGG database.

### Gene set enrichment analysis

Gene set analysis was performed to determine the statistically significant gene sets by GSEA software version 4.1.0 (http://www.gsea-msigdb.org/gsea/index.jsp) [57]. The reference gene sets were ‘hallmark’ and ‘C2.CP.KEGG’ gene set collection (MSigDB. V7.5) and the normalized enrichment score (NES) was calculated. We investigated the gene sets of ‘adherence junction’ for E-cad (CDH1) expression and ‘glycine serine and threonine metabolism’ for serine synthesis pathway.

### Cell culture

MDA-MB-231 and BT549 cells were cultured in Roswell Park Memorial Institute (RPMI) 1640 media (ThermoFisher) and MDA-MB-468 and MCF7 cells were cultured in Dulbecco’s Modified Eagle Medium (DMEM, ThermoFisher) supplemented with 10% (w/v) fetal bovine serum (FBS, ThermoFisher) and 1% (w/v) penicillin/streptomycin solution (ThermoFisher). All cells were maintained at 37 °C in a humidified environment with 5 % CO_2_.

### Drug treatment

Cells were incubated at 37 °C for 6 hr to allow adhesion and spreading onto a cell culture plate before the addition of 15 μM NCT503 (#19718, Cayman) or 10 μM WQ2101 (#SML1970, Millipore Sigma). After 48-hr treatment with the drug, the cells were collected for downstream molecular analyses.

### Cell proliferation and viability assay

For proliferation assay, cells were seeded at a density of 1×10^5^ cells/well onto a 12-well plate and cultured for 48 hr. They were then counted using Countess 3 Cell Counters (ThermoFisher). CellTiter-Glo (#G9242, Promega) was used to compare cell viability after H_2_O_2_ treatment quantitatively.

### Immunoblot analysis

Cells were washed with ice-cold PBS and lysed in RIPA buffer (ThermoFisher) with protease and phosphatase inhibitor cocktail (ThermoFisher). Then, cell lysates were sonicated to shear DNA using a sonicator for 10 to 15 seconds (Branson Sonifier 250, 30% duty cycle, output 4). After centrifuging the lysate at 10,000 g for 5 min at 4 °C, supernatant was collected. Then, protein lysates were separated by SDS-PAGE and transferred to a PVDF membrane, followed by blocking with 5% (w/v) BSA in Tris-Buffered Saline with 0.1% (w/v) Tween 20 (TBST) for 60 min. Then, the membrane was incubated overnight with primary antibodies on a shaker at 4 °C. Membrane was washed three times with TBST for 5 min and then incubated with HRP-conjugated secondary antibodies. After washed three times with TBST for 5 min, ECL Western Blotting Substrate or SuperSignal West Pico PLUS Chemiluminescent Substrate (ThermoFisher) was used for chemiluminescence detection with a Bio-Rad Molecular Imager Gel Doc XR System (Bio-Rad). Primary and secondary antibodies were diluted according to the manufacturer’s recommendations.

### Genetic modulation of E-cad expression level

E-cad perturbations were expressed using the pLKO.1 non-target scramble (#SHC016, Millipore Sigma) and E-cad shRNA (TRCN0000237841, target sequence; AGATTGCACCGGTCGACAAAG, Millipore Sigma). Lentiviral supernatant was prepared by co-transfecting HEK-293T cells with lentivirus packaging plasmids, psPAX2 (Addgene) and pMD2.G (Addgene), and polyethylenimine (1 μg/mL) for 48 hr incubation. Lentivirus-containing media was centrifuged and added to MDA-MB-468 cells for shRNA transduction with Polybrene (8 μg/mL, # sc-134220A, Santa cruz biotechnology). Then, stable cell lines were selected after 1 week of puromycin (2 μg/mL) treatment. The lentiviral E-cadherin-EGFP were generated from pCS-CG (Addgene) for E-cadherin knock-in of MDA-MB-231 as previously described [58]. For siRNA transfection, non-targeting control siRNA and siRNA targeting PHGDH, c-Myc, and SIRT2 (ON-TARGET plus SMART pool siRNA, Dharmacon) were used with Lipofectamine RNAiMAX Transfection Reagent (#13778075, ThermoFisher). Knock-down and knock-in were confirmed by western blot.

### ROS measurement

CellROX green or CellROX orange (ThermoFisher) was applied to live cells cultured on glass bottom dishes (Cellvis) for 30 min according to the manufacturer’s instruction. ROS images were obtained using a Zeiss LSM 780 confocal microscope (Zeiss) with Plan-Apo 20× or 40× Oil lens.

### Immunofluorescence microscopy

Immunofluorescence images were obtained as previously described [59]. Briefly, cells on a glass bottom dish were fixed with paraformaldehyde (PFA) for 10 min at room temperature. Then, fixed cells were permeabilized with 0.1% (w/v) Triton X-100 in PBS for 10 min at room temperature, followed by washing with PBS containing 0.1% (w/v) BSA. Cells were blocked with 5% (w/v) BSA for 30 min. Then, cells were incubated with primary antibodies for 1 hr at room temperature. Nucleus and actin filament were stained with DAPI and DyLight 650 phalloidin (#12956, Cell Signaling Technology) respectively. After washing with PBS containing 0.1% (w/v) BSA, immunofluorescence images were obtained using a Zeiss LSM 780 confocal microscope (Zeiss) with Plan-Apo 20× or 40× Oil lens.

### RNA extraction and qPCR analysis

For quantification of mRNAs, cells were collected, and RNA was extracted using RNeasy Kits (Qiagen) according to the manufacturer’s instructions. Then, qPCR was conducted with iTaq-SYBR Green (Bio-Rad) and primers (Integrated DNA Technologies) using CFX 384 Touch Real-Time PCR detection system (Bio-Rad). We used following primers; E-cad (FWD: TTGCACCGGTCGACAAAGGAC / REV: TGGAGTCCCAGGCGTAGACCAA), c-Myc (FWD: CAGCTACGGAACTCTTGTGC / REV: CAAGACTCAGCCAAGGTTGT), PHGDH (FWD: AACTTCTTCCGCTCCCATTT / REV: GTCATCAACGCAGCTGAGAA), GAPDH (FWD: GCACCGTCAAGGCTGAGAAC / REV: TGGTGAAGACGCCAGTGGA), β-actin (FWD: CGTACCACTGGCATCGTGAT / REV: GTGTTGGCGTACAGGTCTTTG), and HK2 (FWD: CCAGTTCATTCACATCATCAG / REV: CTTACACGAGGTCACATAGC). β-actin mRNA level was used to normalize all the qPCR data.

### Isotope tracing analysis with [U-^13^C] glucose using LC-MS

MDA-MB-231 cells were seeded in a 6-well plate at a density of 5×10^5^ cells/well and allowed to adhere for 12 hr. Then, the culture media was replaced with 2 mL of fresh media containing dialyzed FBS (#26400044, ThermoFisher) and 11 mM [U-^13^C] glucose (#389374, Millipore Sigma). After 6-hr incubation, the cells were washed three times with 0.9% (w/v) NaCl followed by incubation with 500 μL ice-cold extraction solvent (80% methanol/distilled water) (w/v) for 15 min at -80 °C. Then, cells were placed on dry ice and scraped with a cell lifter (#10062-904, VWR). The samples were transferred to a 1.5 mL tube and spun down with 15,000 g for 5 min at 4 °C. The supernatant was transferred to a new 1.5 mL tube. The sample was collected again with 500 μL 80% (w/v) methanol and dried. Metabolites were extracted from the dried pellet by adding 100 μL 40:40:20 acetonitrile:methanol:water. The solvent was vortexed and centrifuged at 16,000 g for 10 min at 4 °C, and 30 μL of the supernatant was transferred into LC-MS vials. A quadrupole orbitrap mass spectrometer (Q Exactive, ThermoFisher) operating in a negative ion mode was coupled to a Vanquish UHPLC system with electrospray ionization and used to scan from m/z 70 to 1,000 at 2 Hz, with a 140,000 resolution. LC separation was achieved with an XBridge BEH Amide column (2.1×150 mm, 2.5 μm particle size, 130 Å pore size; Waters Corporation) using a gradient of solvent A (95:5 water: acetonitrile with 20 mM ammonium acetate and 20 mM ammonium hydroxide, pH 9.45) and solvent B (acetonitrile). The flow rate was 150 μL/min. The LC gradient was: 0 min, 85 % (w/v) B; 2 min, 85 % B; 3 min, 80 % B; 5 min, 80 % B; 6 min, 75 % B; 7 min, 75 % B; 8 min, 70 % B; 9 min, 70 % B; 10 min, 50 % B; 12 min, 50 % B; 13 min, 25 % B; 16 min, 25% B; 18 min, 0% B; 23 min, 0 % B; 24 min, 85 % B; and 30 min, 85 % B. The autosampler temperature was 5 °C and the injection volume was 3 μL. Data were analyzed using the MAVEN software (Build 682, http://maven.princeton.edu/index.php). Natural isotope correction for dual isotopes was performed with AccuCor2 R code (https://github.com/wangyujue23/AccuCor2).

### Lactate generation and glucose consumption analysis

The lactate and glucose levels in culture media were analyzed by commercial kits, Lactate-Glo Assay (#J5021, Promega) and Glucose-Glo Assay (#J6021, Promega), respectively. Each cell line was plated at a concentration of 2×10^5^ cells/mL. After 48-hr incubation, culture media was collected and filtered with Amicon Ultra-0.5 centrifugal filter devices (#UFC500396, Millipore Sigma), and then diluted with PBS (1:300 ratio). The assays were performed according to the manufacturer’s instructions, and luminescence signal levels were measured in a plate reader (Synergy H4 Microplate Reader, Biotek). To calculate the lactate generation or glucose consumption, the differences between the lactate or glucose level after 48-hr incubation and its initial level was normalized to the cell number.

### Oxygen consumption rate (OCR) measurement

Cells were plated on a Seahorse XF-96 (Agilent) plate at a concentration of 1×10^4^ cells/well and allowed to adhere for 6 hr. Culture media was replaced with 172 μL of XF RPMI assay media containing 11 mM XF Glucose, 1 mM XF Pyruvate, and 2 mM XF Glutamine (Agilent), and then cells were incubated in a CO_2_-free incubator for 1 hr. OCR was measured in the presence of 2.5 μM oligomycin, 1.5 μM FCCP, and 0.2 μM rotenone / 2.5 μM antimycin according to manufacturer’s instructions.

### Reduced glutathione (GSH) and oxidized glutathione (GSSG) measurement

GSH and GSSG levels were measured by a commercial kit, GSH/GSSG-Glo Assay (#V6611, Promega), according to the manufacturer’s instructions. Briefly, cells were seeded at a concentration of 1×10^4^ cells/well in a 96-well plate and allowed to adhere for 6 hr. Then, the culture media was replaced with total glutathione lysis reagent or GSSG lysis reagent to lyse cells, followed by shaking the culture plate for 5 min. After adding the luciferin generation reagent, the plate was incubated for 30 min at room temperature. Luminescence was measured after 15-min incubation at room temperature with luciferin detection reagent using a plate reader (Synergy H4 Microplate Reader, Biotek). The GSH/GSSG ratio was calculated using the following formula.

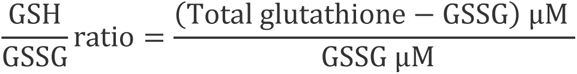

### E-cad functional blocking experiment

MDA-MB-231 E-cad^+^ and MDA-MB-468 E-cad^+^ cells were seeded at a concentration of 2×10^5^ cells/well onto a 6-well plate. Then, 10 μg/mL of E-cadherin functional blocking antibody (HECD-1, #13-1700, ThermoFisher) was added to each well, followed by incubation for 48 hr. Then, cells were observed and collected for immunoblot.

### Histology

For lung isolation from mice, a 20-gauge angiocatheter was sutured into the trachea and the lungs were inflated with 1.5% (w/v) agarose (Boston BioProducts) through the trachea. Once the agarose was solidified after 2 min, the lungs were extracted, and one lobe was fixed with 10 % (w/v) formalin (VWR) for 24 hr. At a Johns Hopkins Medical Institute internal core, samples were paraffin embedded, sectioned, and H&E or immunohistochemistry stained. To extract DNA, the remaining lobes of isolated lungs were flash frozen in liquid nitrogen before storing at -80 °C. Then, the lungs were digested, and DNA was extracted using Genomic DNA mini kit (#K182001, Invitrogen).

### Orthotopic xenograft mouse models

All animal procedures were conducted according to the protocol approved by the Johns Hopkins University Institutional Animal Care and Use Committee. 1×10^6^ cells of E-cad^+^ or E-cad^-^ MDA-1 MB-231 were resuspended in 100 μL of DPBS:Matrigel (1:1 ratio, #354234, Corning) and were injected into the second mammary fat pad of 5-week-old female NSG mice (n = 10). Then, mice were randomly divided into two groups after 7 days, which were assigned blindly to a vehicle or PHGDH inhibitor (NCT503) treatment group. NCT503 solution (#SML1659, Millipore Sigma) was prepared in a 5% (w/v) ethanol, 35% (w/v) polyethylene glycol 300 (#8.17002, Millipore Sigma), and 60%(w/v) aqueous 30% (w/v) (2-Hydroxypropyl)-β-cyclodextrin (#H5784, Millipore Sigma). 40 mg/kg NCT503 was injected intraperitoneally. Tumor volume was measured every 2 days with a caliper and calculated using the formula (0.5 × width^2^ × length). Mice were treated with vehicle or NCT503 solution daily for 20 days.

### 2D migration assay

For 2D single-cell migration, cells were plated on collagen-I-coated tissue culture plastic dishes (20 µg/ml, collagen I rat tail, Thermo Fischer) at 5-8% confluency. After cells spread upon incubation at 37 °C with 5% CO_2_, the culture media was replaced with fresh media with PHGDH inhibitor NCT503 (15 µM). Acquisition of time-lapse images started 30 min after the media change. Cells were imaged in phase contrast mode every 20 min for 20 hr using a Nikon Eclipse Ti inverted microscope (Nikon) equipped with an automated stage control (NIS-Elements; Nikon) and a 10×/0.45 NA objective. During the experiment, a temperature- and CO_2_-controlled stage-top incubator (Tokai Hit) was used to maintain cells at 37 °C with 5% CO_2_. We used the MTrackJ plugin in ImageJ to analyze the cell motility [60] and used a custom MATLAB code (Mathworks) to calculate the cell speed and velocity from the tracking data. For analysis, we chose cells not in contact with the neighboring cells during their motion and stopped the analysis when they contact with another cell. Cells undergoing cell division or apoptosis during the experiment were not included.

### Tail-vein injection and metastasis assay

GFP-tagged breast cancer cells were intravenously injected into mice at a concentration of 10^6^ cells per mouse in a volume of 100 μL. After 48 hr, the lungs were isolated and fixed in formalin for 24 hr. H&E staining and immunohistochemistry were performed to visualize the slides, and lung metastases were imaged using an Axio Scan.Z1 (Zeiss) imaging system. To quantify the metastases, we randomly selected five different sections with an area of 1 mm^2^ per tissue slide and counted anti-GFP signal.

### Statistical analysis

All statistical analyses were performed with GraphPad Prism (GraphPad Software). Significance was assessed with Student’s t test for comparing two groups and one-way analysis of variance (ANOVA) for comparing more than two groups. All experiments in this study were repeated at least three times. Significant differences were summarized in each dataset.

## Results

### E-cad upregulates the *de novo* serine synthesis pathway in breast cancer cells

We hypothesized that E-cad regulates cellular metabolism since E-cad could lead to the hyper-proliferation of breast cancer cells [10], and metabolic reprogramming is a prerequisite for cellular growth and proliferation [16–18]. To identify metabolic genes regulated by E-cad, we performed an unbiased analysis of the proteomics data from tumor pairs xenografted with E-cad knock-in (E-cad^+^) and scramble knock-in (E-cad^-^) MDA-MB-231 breast cancer cells (Fig. 1A) [10]. When we analyzed differentially expressed proteins (DEPs) between E-cad^+^ and E-cad^-^ MDA-MB-231 tumors, we found that multiple metabolic enzymes, including PHGDH and phosphoserine aminotransferase 1 (PSAT1), were upregulated in E-cad^+^ tumors (Fig. 1B).

**Fig. 1.**
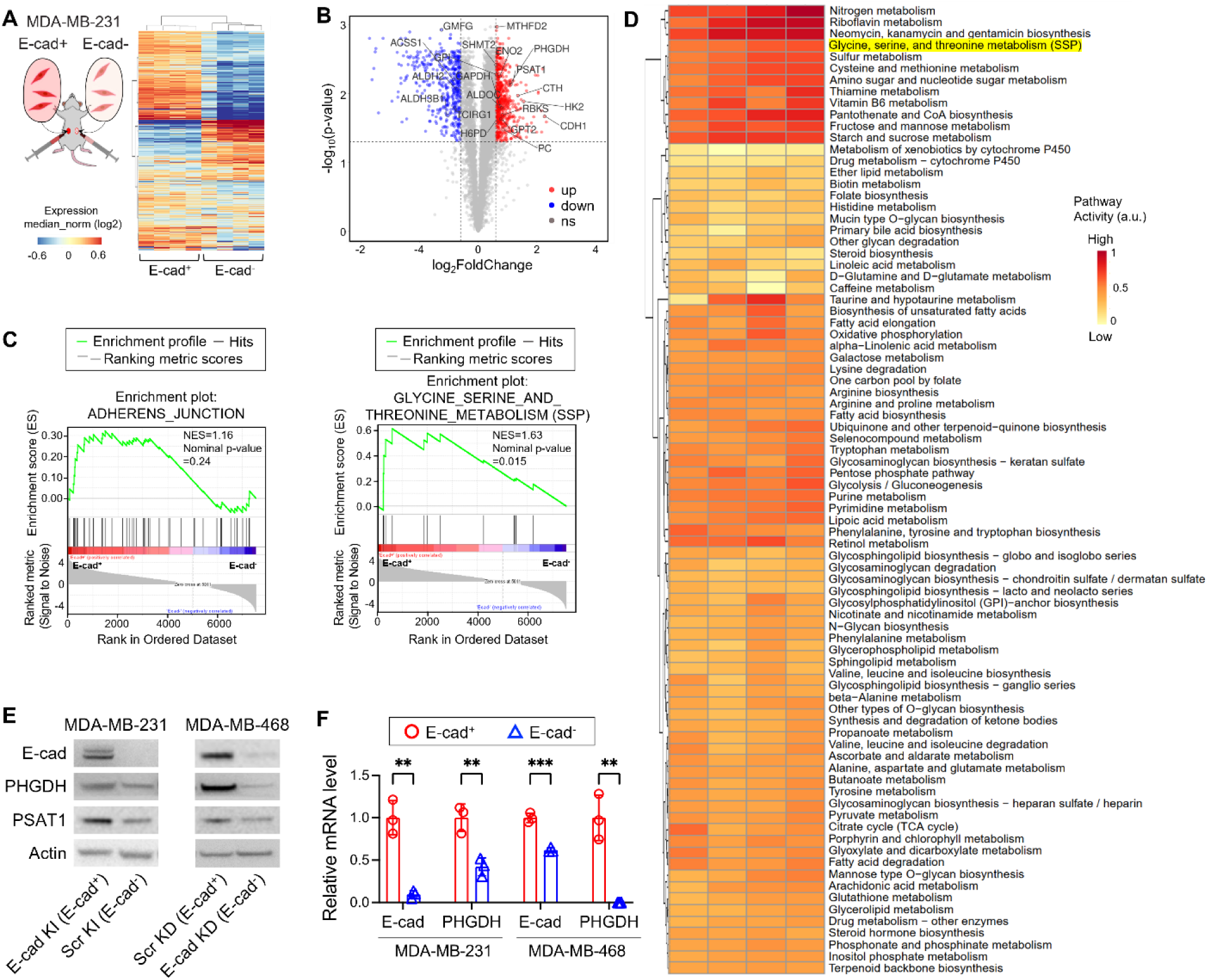
E-cad upregulates the SSP in breast cancer cells. (**A**) Hierarchical clustered heatmap of top 200 differentially expressed proteins generated from proteomics analysis of MDA-MB-231 scramble knock-in (E-cad^-^) and E-cad knock-in (E-cad^+^) bilateral tumors. (**B**) Volcano plot of differentially expressed metabolic enzymes between E-cad^-^ and E-cad^+^ MDA-MB-231 bilateral tumors. Blue and red dots indicate downregulated and upregulated proteins, respectably. ns indicates non-significant (Fold change, ≥ 1.5; q value < 0.05). (**C**) Gene set enrichment analysis (GSEA) demonstrating the enrichment of adherens junction and glycine, serine and threonine metabolism pathways in E-cad^+^ MDA-MB-231 cells. (**D**) Metabolic pathway activities analysis based on differentially expressed proteins between MDA-MB-231 Ecad- and Ecad^+^ bilateral tumors. The protein-coding genes were grouped into 84 metabolic pathways based on KEGG classifications. (**E**) Representative immunoblot analysis of E-cad and SSP enzymes in E-cad^+^ and E-cad^-^ MDA-MB-231 and MDA-MB-468 cells. (**F**) Comparison of mRNA expression of E-cad and PHGDH between E-cad^+^ and E-cad^-^ cells. Statistical analyses were conducted with unpaired two-tailed t test. Error bars indicate SEM (**: p < 0.01; ***: p < 0.001).

PHGDH and PSAT1 are SSP enzymes, and our gene set enrichment analysis (GSEA) on the DEPs confirmed that the SSP was upregulated in E-cad^+^ tumors (NES=1.63, *p*-value < 0.05) (Fig. 1C). For comprehensive analysis of metabolic changes induced by E-cad, we then investigated metabolic pathway variation between E-cad^+^ and E-cad^-^ tumors and quantified the metabolic pathway activities [19]. Among 84 metabolic pathways based on KEGG classifications, the “glycine, serine, and threonine metabolism pathway”, whose gene set includes SSP enzymes, was one of the most upregulated pathways (Fig. 1D and Supplementary Fig. S1A). In addition, principal component analysis (PCA) of the SSP enzyme fold changes showed a clear difference between E-cad^+^ and E-cad^-^ tumors (Supplementary Fig. S1B).

In the bilateral breast cancer xenograft model (Fig. 1A), E-cad^+^ tumors were significantly larger than E-cad^-^ tumors [10]. Therefore, the cells in E-cad^+^ tumors were more likely to face nutrient limitation or hypoxia than those in E-cad^-^ tumors [20]. Given that PHGDH and other SSP enzymes can be upregulated in nutrient-limited or hypoxic conditions [21, 22], the higher expression of PHGDH and PSAT1 in E-cad^+^ tumors might be attributed to environmental factors rather than E-cad itself. To test this hypothesis, we cultured the same clones of MDA-MB-231 cells (E-cad^+^ for E-cad knock-in and E-cad^-^ for scramble knock-in) *in vitro*. Consistent with the *in vivo* proteomics data, protein levels of PHGDH and PSAT1 were significantly higher in E-cad^+^ cells than in E-cad^-^ cells (Fig. 1E and Supplementary Fig. S1C). While MDA-MB-231 cells intrinsically express a low level of E-cad, MDA-MB-468 breast cancer cells express a high level of E-cad (Supplementary Fig. S1D) [23]. E-cad knock-down in MDA-MB-468 cells significantly decreased the expression of PHGDH and PSAT1 (E-cad^+^ for scramble knock-down and E-cad^-^ for E-cad knock-down), further confirming our finding that the SSP enzymes are regulated by E-cad (Fig. 1E). Additionally, we found that the regulation occurred at the transcriptional level (Fig. 1F). Other two breast cancer cell lines, BT549 and MCF7, exhibited a consistent relationship between E-cad and SSP enzymes with E-cad overexpression or knock-down, respectively (Supplementary Fig. S1E). Taken together, our data demonstrate that E-cad positively regulates the SSP in breast cancer cells.

### E-cad^+^ breast cancer cells exhibit higher activities in the SSP and mitochondrial metabolism than E-cad^-^ cells

To determine whether the SSP flux was affected by E-cad, we cultured E-cad^+^ or E-cad^-^ MDA-MB-231 cells in [U-^13^C] glucose media for 6 hr. Liquid chromatography-mass spectrometry (LC-MS) analysis of intracellular metabolites indicated that E-cad^+^ cells exhibited a higher fractional enrichment in serine and glycine than E-cad^-^ cells (Fig. 2A and 2B). Interestingly, the enrichment in tricarboxylic acid (TCA) metabolites, such as succinate and malate, were also higher in E-cad^+^ cells than in E-cad^-^ cells (Fig. 2B). We speculated that the difference in the SSP activity between E-cad^+^ and E-cad^-^ breast cancer cells would be associated with other metabolic pathways, including the TCA cycle, since the SSP is a direct branch of the glycolysis pathway and generates alpha-ketoglutarate (aKG), contributing to TCA anaplerosis [13, 24]. Therefore, we first sought to determine the difference in glucose utilization between E-cad^+^ and E-cad^-^ cells. In both MDA-MB-231 and MDA-MB-458 cell lines, the glucose consumption rate was independent of E-cad level (Fig. 2C). However, in both cell lines, E-cad^+^ cells exhibited a significantly lower generation rate of lactate than E-cad^-^ cells (Fig. 2C). These data indicate that more glucose-derived carbons are used for biosynthetic pathways in E-cad^+^ cells than in E-cad^-^ cells. Also, the oxygen consumption rate (OCR) was significantly higher in E-cad^+^ cells (Fig. 2D and Supplementary Fig. S2A). Multiple studies reported that canonical Wnt signaling pathway, often de-activated by E-cad [25, 26], regulates the expression of monocarboxylate transporter 1 and 4 (MCT1 and MCT4, respectively), which are major lactate transporters, and pyruvate dehydrogenase kinase (PDK), which phosphorylates and inhibits pyruvate dehydrogenase (PDH) (Fig. 2A) [27, 28]. We confirmed that E-cad^+^ cells expressed lower levels of MCT1 and MCT4 and lower phosphorylation of PDH than E-cad^-^ cells (Fig. 2E and Supplementary Fig. S2B), explaining lower lactate generation and higher mitochondrial metabolism in E-cad^+^ cells (Fig. 2B-D). In addition, we found that exogeneous serine uptake was significantly lower in E-cad^+^ cells while the expression level of ASCT2, a primary serine transporter, was marginally different between E-cad^+^ and E-cad^-^ cells (Supplementary Fig. S2C and S2D). Altogether, our metabolic analyses clearly demonstrate that E-cad reprograms cellular metabolism, especially the SSP, in breast cancer cells.

**Fig. 2.**
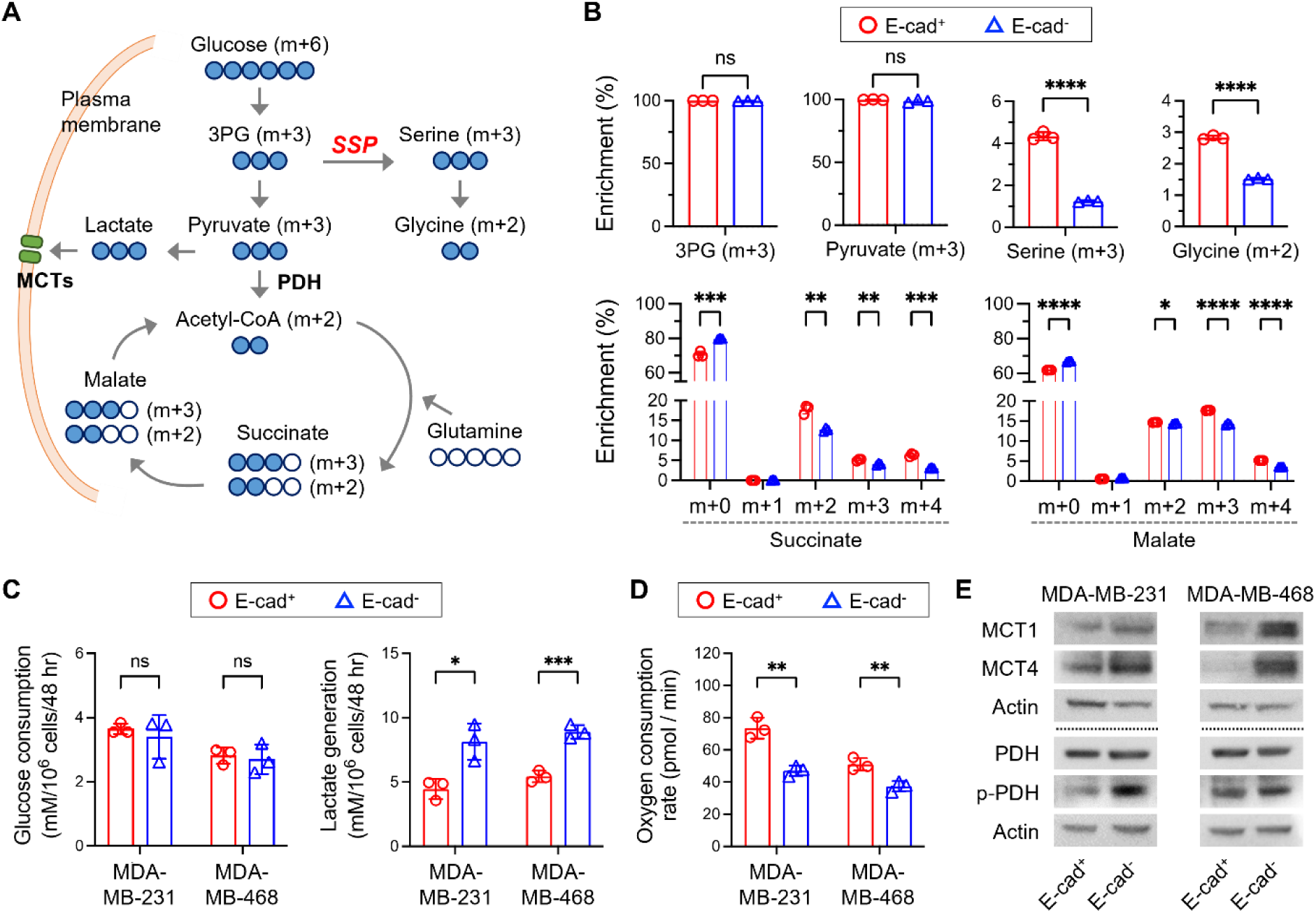
E-cad alters the metabolic activities to SSP from glycolysis. (**A**) Schematic of isotope tracing analysis with [U-^13^C] glucose. 3PG, 3-phosphoglycerate. (**B**) Fractional enrichment in intracellular metabolites in E-cad^+^ and E-cad^-^ MDA-MB-231 cells after 6-hr incubation with [U-^13^C] glucose. (**C**) Glucose consumption and lactate generation rates (mM/10^6^ cells/48 hr) of E-cad^+^ and E-cad^-^ MDA-MB-231 and MDA-MB-468 cells. (**D**) Oxygen consumption rate (OCR) of E-cad^+^ and E-cad^-^ MDA-MB-231 and MDA-MB-468 cells. (**E**) Representative immunoblot analysis of MCT1, MCT4, and PDH in E-cad^+^ and E-cad^-^ MDA-MB-231 and MDA-MB-468 cells. Statistical analyses were conducted with unpaired two-tailed t test. Error bars indicate SEM (ns: not significant; *: p < 0.05; **: p < 0.01; ***: p < 0.001; ****: p < 0.0001).

### The upregulated SSP supports the proliferation of E-cad^+^ breast cancer cells

The SSP provides precursors for biosynthetic pathways, and its inhibition hampers the proliferation of cancer cells [14, 29, 30]. Since E-cad was shown to accelerate the proliferation of cancer cells [7, 10, 31], we hypothesized that higher proliferation of E-cad^+^ breast cancer cells could be attributed to their upregulated SSP. To test this hypothesis, we compared the growth rates of E-cad^+^ and E-cad^-^ MDA-MB-231 and MDA-MB-468 cells with PHGDH inhibition.

While E-cad^+^ cells proliferated much faster than E-cad^-^ cells, PHGDH inhibitor treatment (NCT503 or WQ2101) or PHGDH knock-down via siRNA significantly and specifically hindered the proliferation of E-cad^+^ cells (Fig. 3A and 3B and Supplementary Fig. S3A). The accelerated proliferation was also observed in other breast cancer cell lines, BT549 and MCF7 (Supplementary Fig. S3B).

**Fig. 3.**
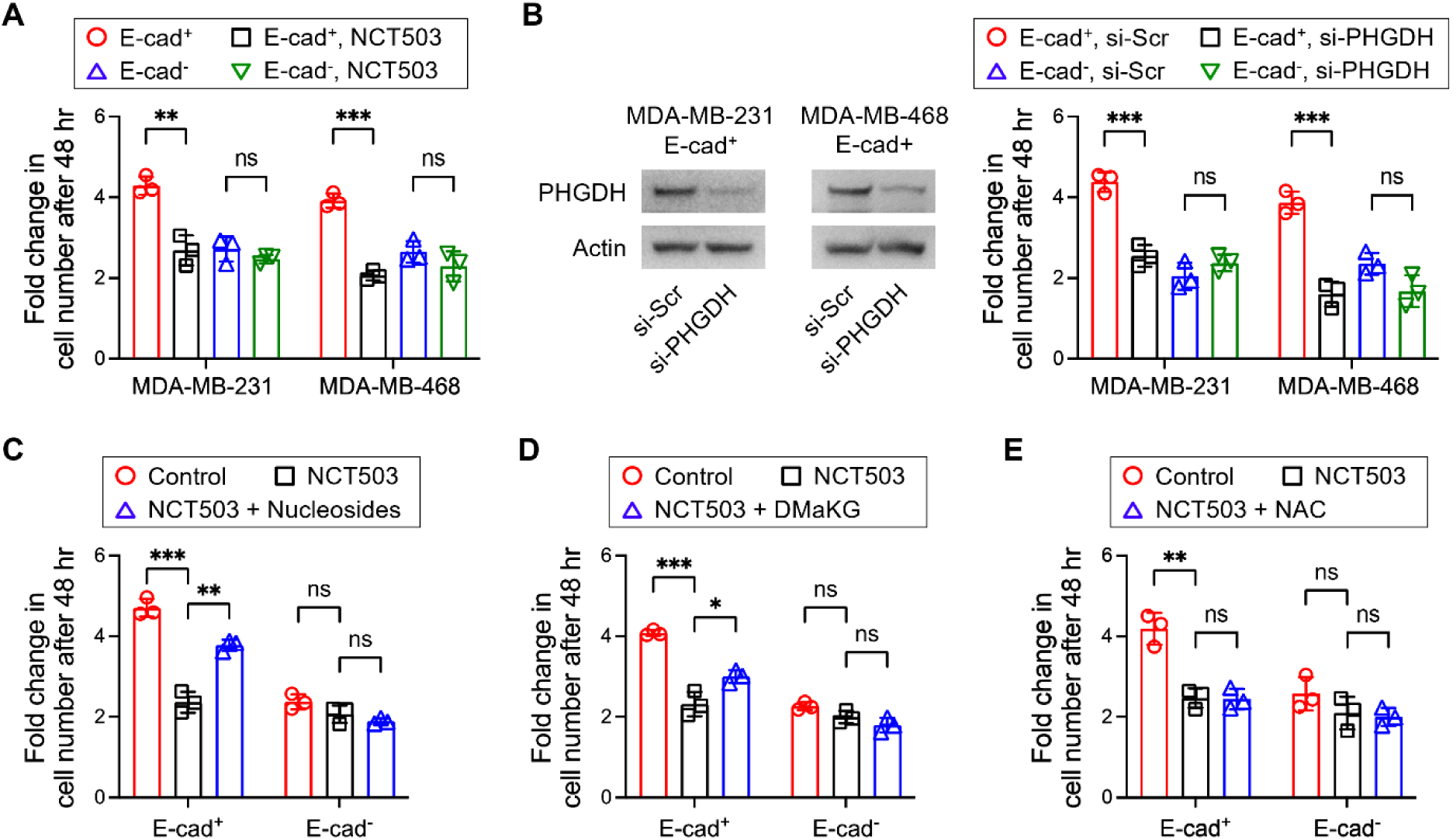
The upregulated SSP promotes the biosynthesis in E-cad^+^ cancer cells, consequently promoting their proliferation. (**A**) Fold change in cell number of E-cad^+^ and E-cad^-^ MDA-MB-231 and MDA-MB-468 cells with PHGDH inhibition by NCT503. (**B**) Fold change in cell number of E-cad^+^ and E-cad^-^ MDA-MB-231 and MDA-MB-468 cells with PHGDH inhibition by siRNA. (**C**) Fold change in cell number of E-cad^+^ MDA-MB-231 cells with NCT503 and nucleoside supplementation. (**D**) Fold change in cell number of E-cad^+^ MDA-MB-231 cell with NCT503 and dimethyl-alpha-ketoglutarate (DMaKG). (**E**) Fold change in cell number of MDA-MD-231 cells with NCT503 and ROS scavenger, NAC. Statistical analyses were conducted with unpaired two-tailed t test. Error bars indicate SEM (ns: not significant; *: p < 0.05; **: p < 0.01; ***: p < 0.001).

To identify the underlying mechanism of how the SSP affects proliferation of E-cad^+^ breast cancer cells, we investigated three major roles of the SSP: nucleotide synthesis [32], aKG generation [13], and ROS management [33]. First, we tested whether the effect of SSP inhibition could be reversed by supplementation of nucleosides (cytidine, guanosine, uridine, adenosine, and thymidine) or dimethyl-aKG (cell-permeable form of aKG). Indeed, the reduced proliferation of E-cad^+^ MDA-MB-231 cells with NCT503 treatment was significantly rescued with nucleosides or dimethyl-aKG (Fig. 3C and 3D). Interestingly, NCT503 treatment also specifically elevated the ROS level in E-cad^+^ cells (Supplementary Fig. S3C and S3D), but the supplementation of ROS scavenger, N-acetylcysteine (NAC), barely rescued their proliferation while reducing the ROS level (Fig. 3E and Supplementary Fig. S3E and S3F). These data indicate that the upregulated SSP in E-cad^+^ breast cancer cells promotes nucleotide synthesis and TCA anaplerosis, clear indication of their metabolic advantage for proliferation compared to E-cad^-^ breast cancer cells [10].

### The upregulated SSP makes E-cad^+^ cells resistant to oxidative stress

While the ROS scavenger could not rescue the proliferation of E-cad^+^ cells treated with PHGDH inhibitor, the basal ROS levels were lower in E-cad^+^ cells than in E-cad^-^ cells (Supplementary Fig. S3C and S3D). Since the SSP provides precursors for the synthesis of glutathione, a major antioxidant, we hypothesized that E-cad^+^ breast cancer cells could have a higher capacity to reduce oxidative stress than E-cad^-^ cells. To investigate this hypothesis, we induced oxidative stress in E-cad^+^ or E-cad^-^ breast cancer cells with 100 μM hydrogen peroxide (H_2_O_2_) for 30 min. Under this acute exogenous oxidative stress, E-cad^+^ MDA-MB-231 and MDA-MB-468 cells exhibited markedly lower levels of ROS than E-cad^-^ counterparts, but their oxidative stress resistance was reduced with PHGDH knock-down (Fig. 4A and 4B). The differences in ROS-reducing capacity were reflected in the differences in cell viability after 2-hr incubation with H_2_O_2_: E-cad^+^ cells survived significantly more than E-cad^-^ cells, but their survival advantage was diminished with PHGDH knock-down (Fig. 4C). As expected, the ratio of reduced to oxidized glutathione (GSH/GSSG) was significantly higher in E-cad^+^ cells than E-cad^-^ cells, and PHGDH knock-down in E-cad^+^ cells reduced the ratio (Fig. 4D). These results demonstrated that the upregulated SSP plays a vital role in E-cad^+^ cells on mitigating oxidative stress.

**Fig. 4.**
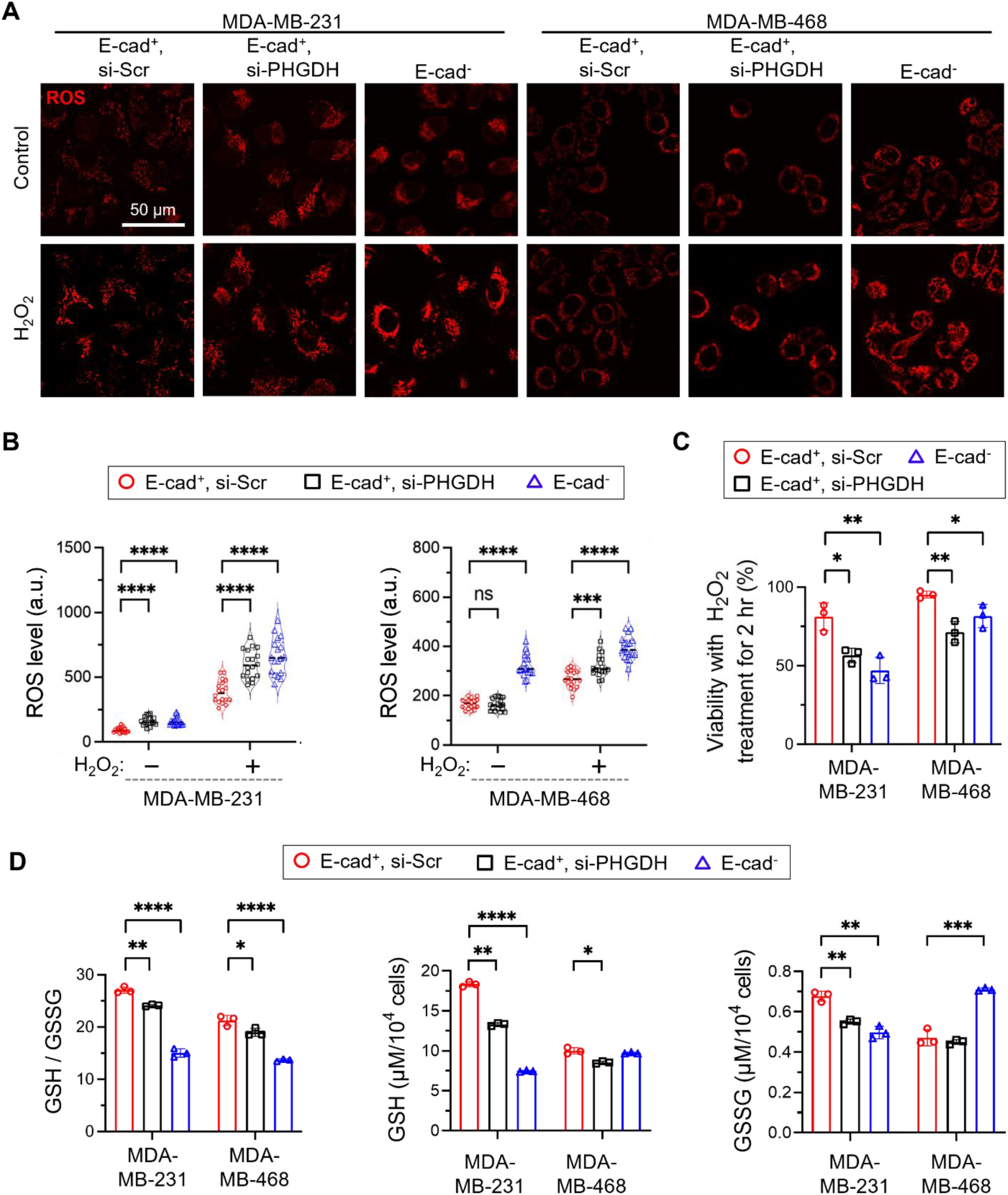
SSP upregulation enhances oxidative stress resistance of E-cad^+^ cancer cells. (**A**) Representative fluorescence images of intracellular ROS in E-cad^+^ and E-cad^-^ cells cultured in the presence or absence of 100 μM H_2_O_2_ for 30 min. (**B**) Quantified levels of ROS, indicating higher resistance against exogenous oxidative stress in E-cad^+^ cells (n = 18). a.u., arbitrary units (**C**) Cell viabilities in the presence of 100 μM H_2_O_2_ for 2 hr. (**D**) Glutathione (GSH) and oxidized glutathione (GSSG) ratio in E-cad^+^ and E-cad^-^ cells. Statistical analyses were conducted with unpaired two-tailed t test. Error bars indicate SEM (ns: not significant; *: p < 0.05; **: p < 0.01; ***: p < 0.001; ****: p < 0.0001).

### SIRT2-mediated E-cad downregulation reduces the PHGDH expression and diminishes the ROS-reducing capacity

We then tested whether EMT induction which downregulates E-cad could also affect the PHGDH expression level in breast cancer cells. Sirtuin 2 (SIRT2), a NAD^+^-dependent protein deacetylase, was reported to suppress the E-cad expression and induce EMT in breast cancer cells [34, 35]. Therefore, we hypothesized that SIRT2 could negatively regulate PHGDH expression. Indeed, SIRT2 overexpression (OV) in E-cad^+^ MDA-MB-231 and MDA-MB-468 cells decreased the expression levels of E-cad and PHGDH (Fig. 5A and 5B) and cellular proliferation (Fig. 5C). This is consistent with our findings that E-cad^+^ cells proliferate faster than E-cad^-^ cells due to higher expression of PHGDH (Fig. 3A, 3B, and Supplementary Fig. S3A). To investigate whether SIRT2 OV diminishes ROS-reducing capacity, we treated E-cad^+^ MDA-MB-231 and MDA-MB-468 cells with 100 μM H_2_O_2._ In both cell lines, SIRT2 OV significantly increased the ROS levels and rendered the cells more vulnerable to H_2_O_2_-induced oxidative stress (Fig. 5D-F); less cells were viable with SIRT2 OV after 2-hr treatment of H_2_O_2_. Also, SIRT2 knock-down in E-cad^-^ MDA-MB-231 and MDA-MB-468 cells showed the opposite results (Supplementary Fig. S4); briefly, SIRT2 knock-down increased the expression levels of E-cad and PHGDH and cellular proliferation, and enhanced the ROS reducing capacity. These results consistently demonstrated that E-cad-mediated PHGDH upregulation helps E-cad^+^ breast cancer cells resist oxidative stress.

**Fig. 5.**
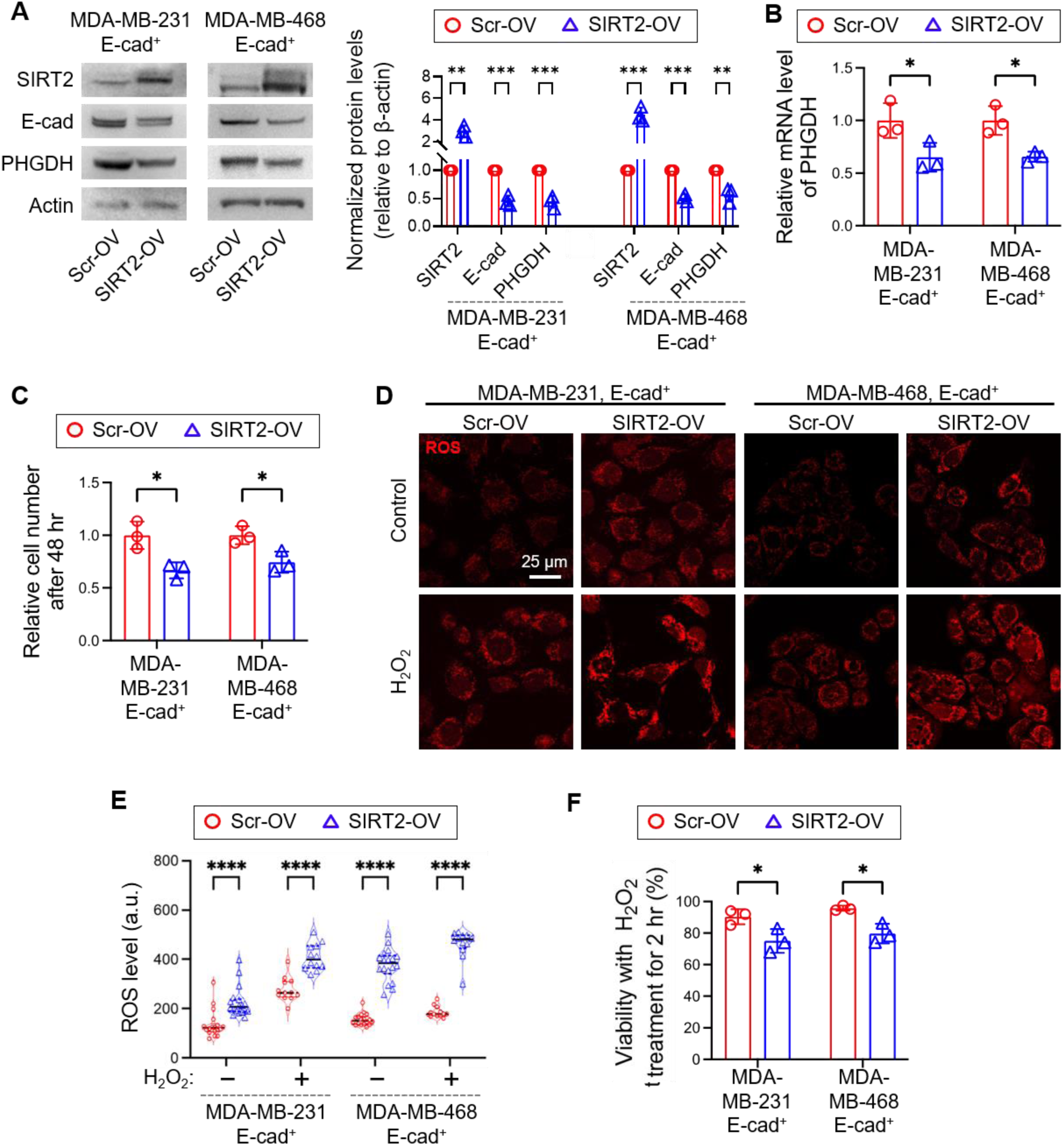
SIRT2-mediated E-cad downregulation decreases PHGDH expression. (**A**) Immunoblot analysis of SIRT2, E-cad, and PHGDH in scramble (Scr)-overexpression (OV) and SIRT2-OV E-cad^+^ cells. (**B**) Comparison of mRNA expression of PHGDH between Scr-OV and SIRT2-OV E-cad^+^ cells. (**C**) Relative cell number of Scr-OV and SIRT2-OV E-cad^+^ cells after 48 hr. (**D**) Representative fluorescence images of intracellular ROS in Scr-OV and SIRT2-OV cells cultured in the presence or absence of 100 μM H_2_O_2_ for 30 min. (**E**) ROS levels in Scr-OV and SIRT2-OV cells cultured in the presence or absence of 100 μM H_2_O_2_ for 30 min. a.u., arbitrary units, n = 18. (**F**) Cell viability in the presence of 100 μM H_2_O_2_ for 2 hr. Statistical analyses were conducted with unpaired two-tailed t test. Error bars indicate SEM (ns: not significant; *: p < 0.05; **: p < 0.01; ***: p < 0.001; ****: p < 0.0001).

### Physical intercellular interaction is not necessary for E-cad-mediated PHGDH upregulation

Since E-cad organizes cell-cell adherens junctions, we investigated whether physical intercellular interaction is required for the PHGDH upregulation. We inhibited the formation of adherens junctions with an E-cad blocking antibody, HECD-1 (10 μg/mL) [36], or a Ca^2+^ chelator, BAPTA (50 μM), because adherens junction formation by E-cad depends on the extracellular Ca^2+^ level [37]. 48-hr treatment of the reagents changed the cell morphology, indicating the disturbance of physical intercellular interactions, but PHGDH expression levels remained unaltered (Supplementary Fig. S5). This implies that juxtacrine signaling from extracellular E-cad domains does not contribute to E-cad-mediated PHGDH upregulation in E-cad^+^ breast cancer cells.

### E-cad upregulates the SSP enzyme expression via c-Myc

To identify the molecular link between E-cad and SSP enzymes, we assessed the expression level of multiple transcriptional factors associated with SSP enzymes: c-Myc [38], activation transcription factor 4 (ATF4) [39], and hypoxia inducible factor-1/2α (HIF-1α and HIF-2α) [40]. Interestingly, the level of c-Myc was significantly higher in E-cad^+^ MDA-MB-231 and MDA-MB-468 cells compared to their E-cad^-^ counterparts. On the other hand, the level of ATF4 was similar between E-cad^+^ and E-cad^-^ cells, while HIF-1α and HIF-2α showed an inconsistent trend between the two groups (Fig. 6A and 6B and Supplementary Fig. S6A). c-Myc knock-down decreased PHGDH and PSAT1 expression in E-cad^+^ cells (Fig. 6C and Supplementary Fig. S6B) and significantly reduced cellular proliferation (Fig. 6D). Overall, these data indicate that the SSP upregulation via c-Myc contributes to the proliferation of E-cad^+^ breast cancer cells. Then, we asked how E-cad regulates c-Myc expression. Consistent with previous work [7], we found that E-cad^+^ MDA-MB-231 cells exhibited lower nucleus-to-cytoplasmic ratios of Smad2/3 than E-cad^-^ cells (Fig. 6E). Smad2 and Smad3 are signaling molecules downstream of TGF-β pathway, and their phosphorylation, followed by its nuclear localization, was shown to suppress the c-Myc expression [41]. Therefore, we reason that the E-cad^+^ cells, with less nuclear localization of Smad2/3, express a higher level of c-Myc than the E-cad^-^ cells.

**Fig. 6.**
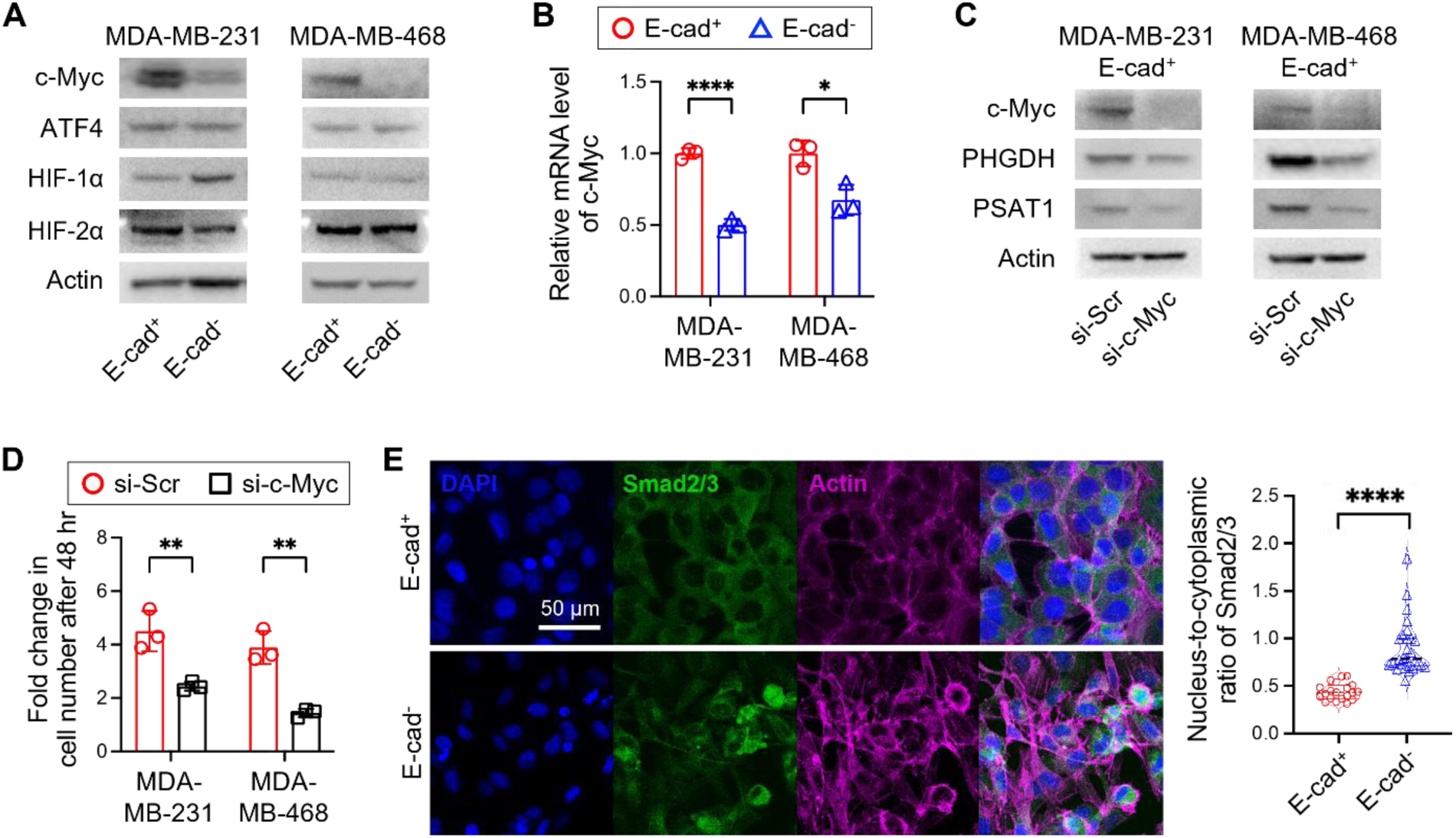
c-Myc mediates the SSP upregulation in E-cad^+^ breast cancer cells. (**A**) Representative immunoblot analysis of c-Myc, ATF4, HIF-1α, and HIF-2α in E-cad^+^ and E-cad^-^ MDA-MB-231 and MDA-MB-468 cells. (**B**) Comparison of mRNA expression of c-Myc between E-cad^+^ and Ecad^-^ MDA-MB-231 and MDA-MB-468 cells. (**C**) Representative immunoblot analysis of c-Myc, PHGDH, and PSAT1 in si-Scr and si-c-Myc E-cad^+^ MDA-MB-231 and MDA-MB-468 cells. (**D**) Fold change in cell number of E-cad^+^ MDA-MB-231 and MDA-MB-468 cells with c-Myc inhibition by siRNA. (**E**) Spatial analysis of Smad2/3 expression in E-cad^+^ and E-cad^-^ cells. Statistical analyses were conducted with unpaired two-tailed t test. Error bars indicate SEM (**: p < 0.01; ****: p < 0.0001).

### PHGDH inhibitor treatment hampers the proliferation and metastasis of E-cad^+^ breast cancer cells *in vivo*

To further corroborate our conclusion that E-cad-mediated SSP upregulation is critically beneficial for proliferation and ROS management, we performed *in vivo* experiments using orthotopic xenograft mouse models. E-cad^+^ and E-cad^-^ MDA-MB-231 cells were implanted into mammary fat pads of NSG mice bilaterally, and when the tumors became palpable (day 8 post-implantation), the mice were daily treated with intraperitoneal injection of NCT503 (Fig. 7A). As shown in other studies [29], NCT503 treatment regimen was well tolerated without noticeable weight loss or physical status changes over the treatment period (Supplementary Fig. S7A). E-cad^+^ tumors grew much slower in NCT503-treated mice than in vehicle-treated mice (Fig. 7B and 7C). In contrast, E-cad^-^ tumors grew much slower than E-cad^+^ tumors and their volumes were similar between NCT503-treated and vehicle-treated mice (Fig. 7B and 7C). This confirmed that the SSP plays a significant role in the proliferation of E-cad^+^ breast cancer cells *in vivo*.

**Fig. 7.**
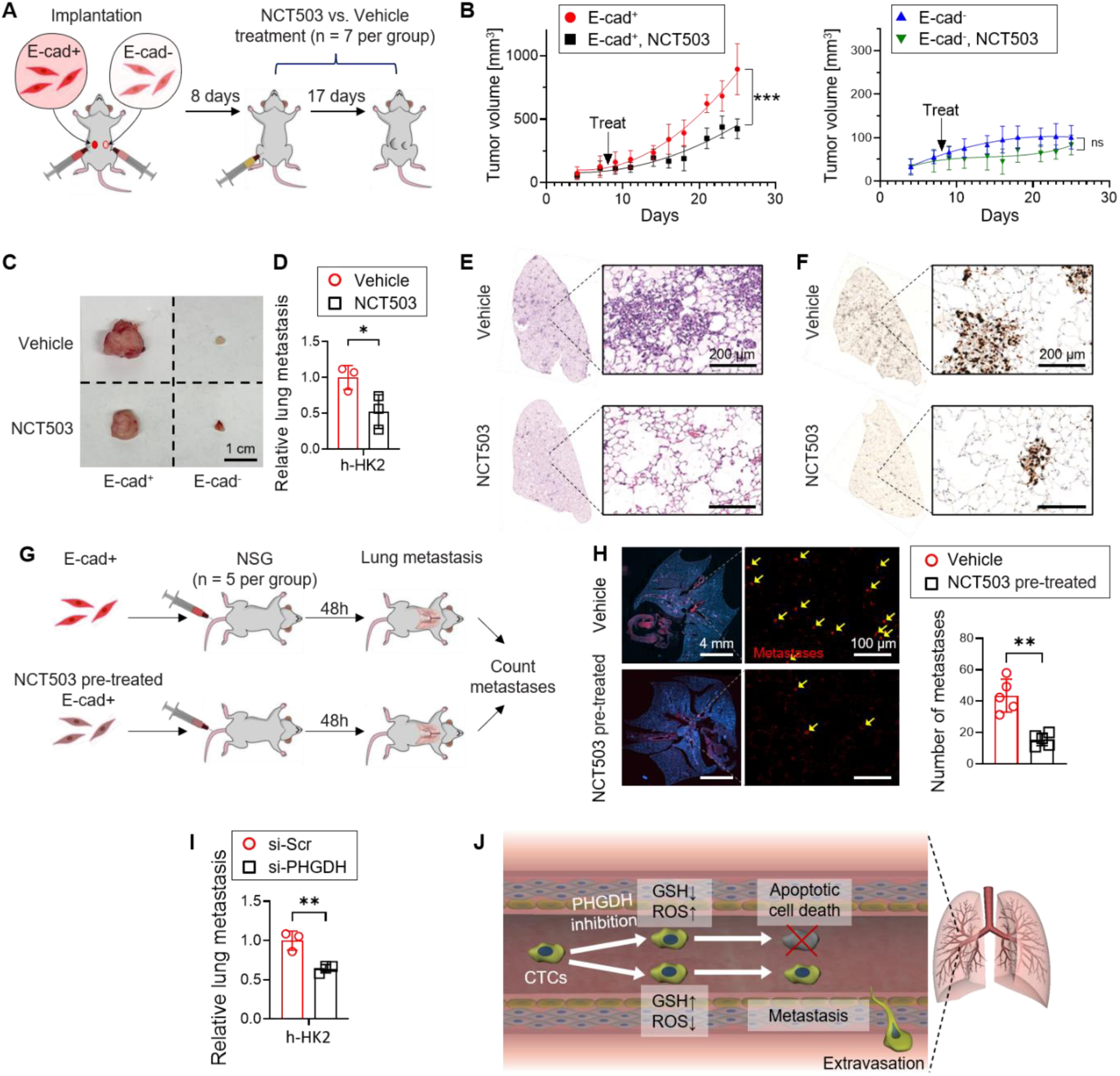
PHGDH inhibitor treatment hampers the proliferation and metastasis of E-cad^+^ breast cancers, not E-cad^-^ breast cancers. (**A**) Schematic diagram of orthotopic implantation of E-cad^+^ and E-cad^-^ MDA-MB-231 cells into mammary fat pad of NSG mice (n = 7). (**B**) Growth curves of E-cad^+^ and E-cad^-^ MDA-MB-231 tumors with vehicle or NCT503 treatment. (**C**) Representative tumor images at the end of the study. (**D**) Quantification of lung metastasis by analyzing human genomic marker, HK2 (hexokinase 2), in the lungs using qPCR. (**E**) H&E staining of the lungs from mice with vehicle or NCT503 treatment. (**F**) Ki67 staining of the lungs from mice with vehicle or NCT503 treatment. (**G**) Schematic diagram of intravenous (IV) injection of E-cad^+^ MDA-MB-231 cells and lung metastasis analysis. (**H**) Comparison of lung metastasis quantified by E-cad^+^ MDA-MB-231 cell number in the lungs (n = 5 mice). Metastases were counted per 1 mm^2^ of lungs. (**I**) Quantification of lung metastasis 48 hr after IV injection of scramble-knockdown (KD) and PHGDH-KD MDA-MB-231 E-cad^+^ cells. (**J**) Schematic diagram of metastasis affected by PHGDH inhibition. Statistical analyses were conducted with unpaired two-tailed t test. Error bars indicate SEM (ns: not significant; *: p < 0.05; **: p < 0.01).

We next sought to determine the effect of PHGDH inhibition on metastasis. While the number of lung metastases in NCT503-treated mice was significantly lower than in vehicle-treated mice (Fig. 7D-F and Supplementary Fig. S7B), this difference could be attributed to the larger size of the primary tumors in the vehicle group (Fig. 7B and 7C), since more cancer cells could be disseminated from larger tumors. Given that ROS could promote epithelial cell migration [42–44], we hypothesized that inhibiting PHGDH in E-cad^+^ breast cancer cells could increase their cell migration due to elevated ROS levels (Supplementary Fig. S3B and S3C). Using our two-dimensional (2D) cell tracking analysis, we confirmed that NCT503 pre-treatment or PHGDH knock-down significantly increased the migration of E-cad^+^ cells, which was reversed by NAC supplementation (Supplementary Fig. S8). To determine whether the enhanced migration with NCT503 pre-treatment leads to more metastases *in vivo*, we injected vehicle- or NCT503-pre-treated E-cad^+^ MDA-MB-231 cells into the tail vein of NSG mice and quantified the lung micro-metastases after 48 hr (Fig. 7G). Interestingly, the micro-metastases were significantly lower in mice injected with NCT503-pre-treated E-cad^+^ cells than in mice with vehicle-pre-treated E-cad^+^ cells (Fig. 7H). Furthermore, we modeled *in vivo* circulation with the constant rotation of media using a magnetic stir bar (250 rotations per minute) in a non-adhesive substrate and assessed the viability of E-cad^+^ or E-cad^-^ MDA-MB-231 cells in this culture system (Supplementary Fig. S9). Consistently, E-cad^+^ cells exhibited higher viability than E-cad^-^ cells after 24-hr culture with the rotation, and their survival advantage was diminished with NCT503 pre-treatment (Supplementary Fig. S8). Previous studies reported that ROS management is critical for circulating tumor cells to resist oxidative stress and survive for metastasis formation [7, 45]. Since the upregulated SSP confers a higher ROS-reducing capacity to E-cad^+^ breast cancer cells (Fig. 4), we reason that it promotes more metastases of E-cad^+^ cells than E-cad^-^ cells. Collectively, these data indicate that the SSP inhibition in E-cad^+^ breast cancer cells increases their migration but hampers their survival during circulation, overall reducing their metastases.

## Discussion

Recent findings suggest that the acquisition of mesenchymal phenotype of epithelial cancer cells, known as EMT, requires distinct metabolic features to comply with the metabolic demands of upregulated metastatic potential [46, 47]. E-cad is a calcium-dependent transmembrane glycoprotein that acts as a glue between cells, and its loss has been associated with EMT, cancer invasion, and metastasis [48, 49]. However, the paradigm that E-cad is a tumor suppressor has been recently challenged since several studies have shown that E-cad indeed supports the proliferation and metastasis of breast cancer cells [7, 10]. While extensive studies have been conducted to identify the role of E-cad in cancer, how E-cad is linked to cancer metabolism has been rarely discussed. Building blocks for tumor growth and ROS-reducing agents for survival during metastasis are generated by metabolic reactions, and therefore metabolic changes associated with E-cad could play a significant role in cancer. Our proteomic and metabolomic analyses of breast cancer cells revealed that E-cad markedly reprograms metabolism, specifically upregulating the SSP, which is critically beneficial for higher proliferation and more metastases of E-cad^+^ breast cancer cells than E-cad^-^ cells.

We also confirmed E-cad-mediated PHGDH upregulation with SIRT2, a transcriptional repressor of E-cad. SIRT2 induces EMT by stabilizes SLUG, which suppresses E-cad expression via AKT/GSK/β-catenin signaling pathway [35]. SIRT2 OV decreased the expression of E-cad and PHGDH in E-cad^+^ breast cancer cells and hampered their proliferation and ROS-reducing capacity. Furthermore, we identified that c-Myc, a major transcription factor regulating metabolism, plays a key role in E-cad-mediated PHGDH upregulation. The SSP enzymes could be regulated by multiple transcription factors, among which the c-Myc expression was consistently elevated in E-cad^+^ breast cancer cells. As expected, c-Myc knock-down decreased the PHGDH and PSAT expression levels as well as the proliferation rate of E-cad^+^ breast cancer cells.

It was previously shown that loss of PHGDH protein in breast cancer cells was associated with elevated invasion, mainly due to integrin α_v_β_3_ sialyation, which enhanced cancer dissemination and metastasis [50]. It is consistent with our findings that E-cad^-^ cells, which expressed lower PHGDH, exhibited a higher migratory propensity than E-cad^+^ cells bearing higher PHGDH. However, the role of PHGDH may vary depending on cell types; its inhibition significantly hampered or rarely affected tumor growth in different cancer models [14, 29, 50–52] or it can be translocated into the nucleus and facilitate malate oxidation in certain cancer cells [53]. Our experiments revealed that E-cad-mediated metabolic reprogramming, especially SSP upregulation, supports tumor growth and metastasis of breast cancers.

In summary, our study revealed that E-cad upregulates the SSP enzymes, including PHGDH, in breast cancers, which is critically beneficial for the growth and metastasis of E-cad^+^ breast cancers. In addition, PHGDH inhibition abrogates the metabolic advantages conferred by E-cad in breast cancer cells, significantly hampering their metastasis. Our findings highlight the role of E-cad-mediated metabolic reprogramming in E-cad^+^ breast cancers.

## Supporting information

https://www.biorxiv.org/content/10.1101/2023.05.24.541452v1.abstract

## Author Contributions

G.L. conducted all biochemical experiments and analyses. C.W. and A.C. assisted *in vitro* experiments and performed *in vivo* experiments. J.J.W. and A.J.E. assisted in designing metastasis experiments and discussed the results. A.J.C. assisted *in vivo* experiments and lung metastasis analysis. G.C.R and D.W. provided cells and proteomics data and discussed the analysis. J.K. and M.J. assisted *in vivo* experiments. L.H. and C.J. performed the LC-MS experiments and analyzed the data. R.D.L. assisted the OCR experiments. B.R.S. and K.K. assisted in designing the migration assay and discussed the results. G.L. and S.J. wrote the manuscript with useful input from all authors.

## Acknowledgement

We thank Dr. Chi V. Dang for providing insightful feedback on our experiments and analyses. This research was supported in part by NIH R00CA226357 (S.J.), P30CA006973 (Sidney Kimmel Comprehensive Cancer Center), U54CA143868 (D.W.), U54CA268083 (D.W.), U01AG060903 (D.W.), U54AR081774 (D.W.), U54CA268083 (A.J.E.), the Breast Cancer Research Foundation BCRF-22-048 (A.J.E.), METAvivor (A.J.E.), the JKTG Foundation (A.J.E.), NIH R01GM142175 (K.K.), R01CA226765 (R.D.L), K99HL150628 (M.J.), Maryland Stem Cell Research Fund Launch Award 2023-MSCRFL-5999 (M.J.), AASLD Foundation Pinnacle Research Award in Liver Disease (C.J.), the Edward Mallinckrodt, Jr. Foundation Award (C.J.), and NIH R01AA029124 (C.J.).

## Data Availability

All the data supporting the findings of this study are available when requested.

