## Supplementary material for "Serine synthesis pathway upregulated by E-cadherin is essential for the proliferation and metastasis of breast cancers": https://www.biorxiv.org/content/10.1101/2023.05.24.541452v1.abstract

### Extended Data Figures

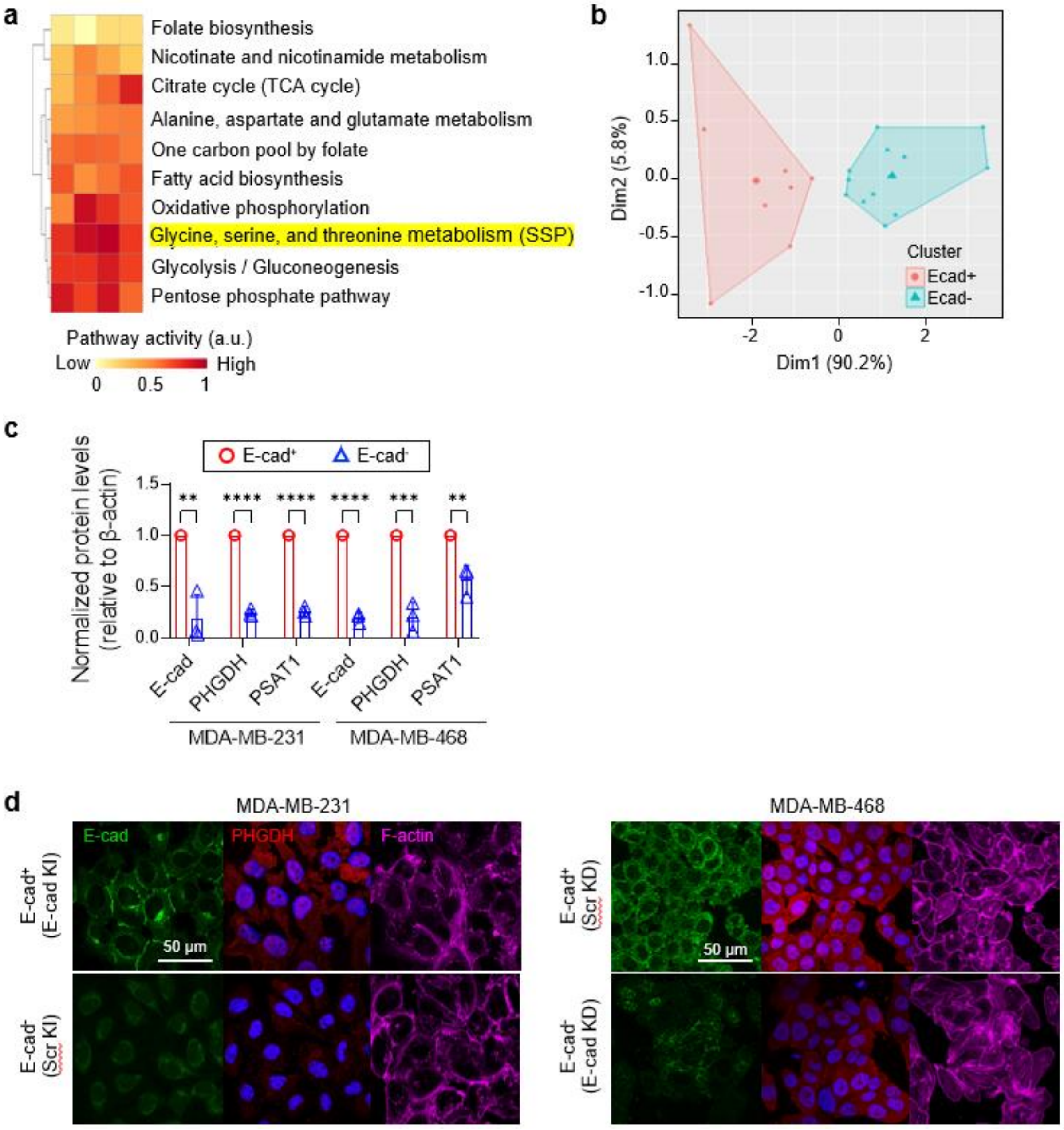


**Extended Data Fig. 1. E-cad upregulates the PHGDH expression in breast cancer cells**
(**a**) Relative pathway activity of top 10 major energy providing metabolic pathways. (**b**) Principal-component analysis of differential expression genes of serine synthesis pathway. The lists of genes were sorted based on KEGG classification. (**c**) Normalized protein levels of E-cad and PHGDH. (**d**) Representative immunofluorescence images of E-cad and PHGDH in E-cad$^{+}$ and E-cad$^{-}$ cells. Expression of PHGDH (red), E-cad (green), and actin filament (Dylight 650) were visualized in nuclear DAPI (blue)-stained breast cancer cells. Statistical analyses were conducted with unpaired two-tailed t test. Error bars indicate SEM (ns: not significant; **: $p < 0.01$; ***: $p < 0.001$; ****: $p < 0.0001$).

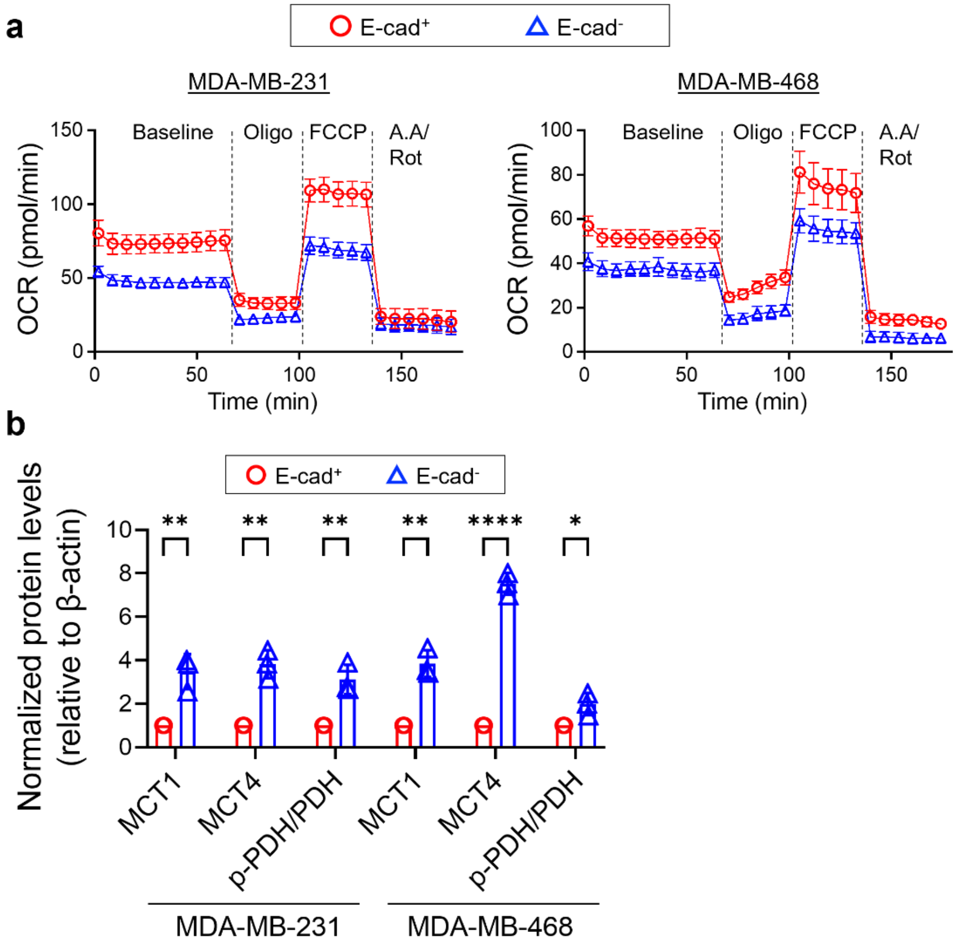


**Extended Data Fig. 2. E-cad$^+$ breast cancer cells exhibit a higher oxygen consumption rate (OCR) than E-cad$^-$ cells.**

(**a**) OCR in response to oligomycin (2 µM), FCCP (0.1 µM), and antimycin A (1 µM) / rotenone (1 µM). (**b**) Normalized protein levels of MCT1, MCT4, and PDH. Statistical analyses were conducted with unpaired two-tailed t test. Error bars indicate SEM (ns: not significant; *: $p < 0.05$; **: $p < 0.01$; ***: $p < 0.001$; ****: $p < 0.0001$).

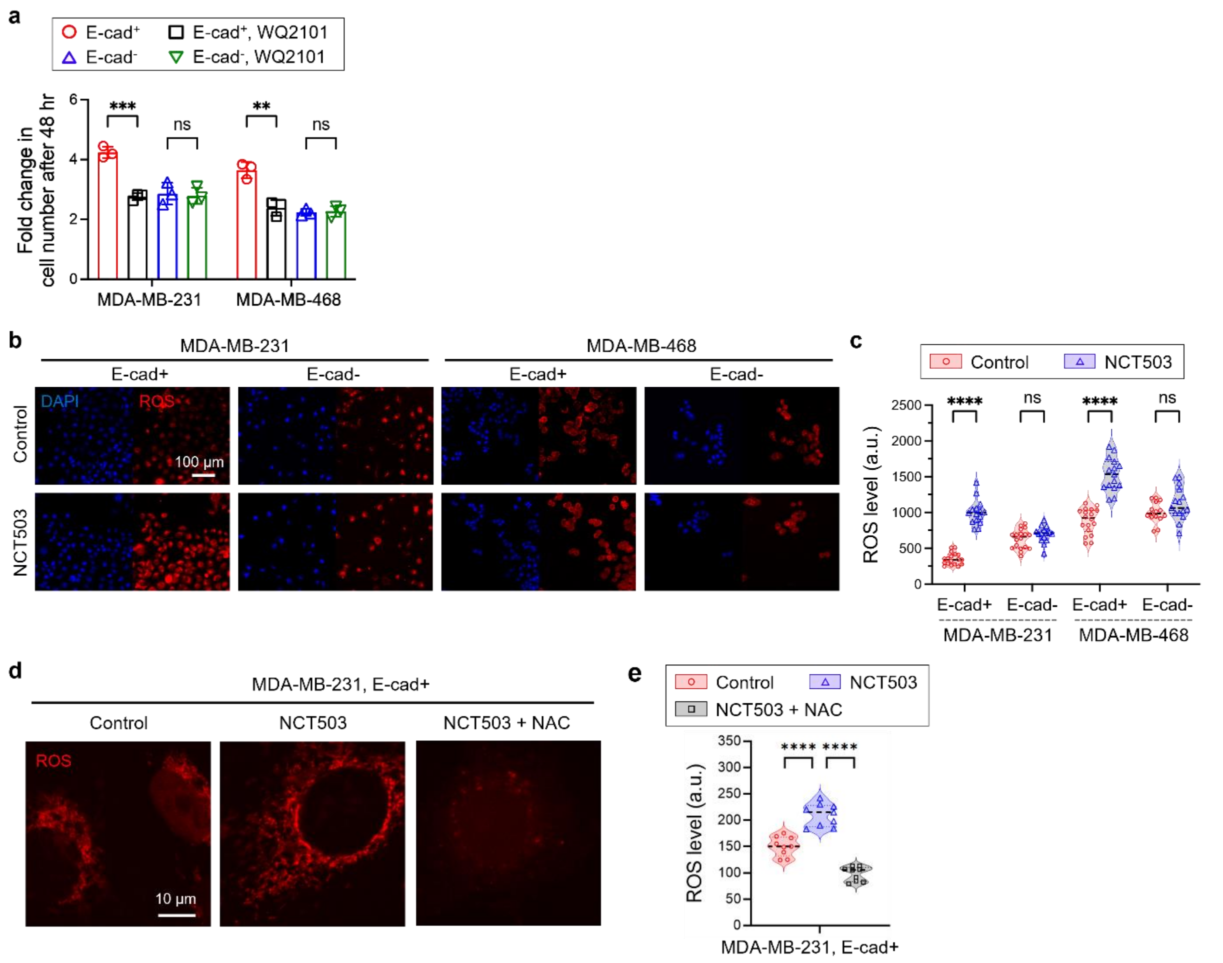


**Extended Data Fig. 3. PHGDH inhibition hampers the proliferation of E-cad$^+$ breast cancer cells and elevates their ROS levels.**

(**a**) Fold change in cell number of breast cancer cells with PHGDH inhibition by 10 μM WQ2101. (**b**) Representative fluorescence images of intracellular ROS in E-cad$^+$ and E-cad$^-$ cells cultured in the presence or absence of 15 μM NCT503 for 48 h. (**c**) Quantified levels of ROS in E-cad$^+$ and E-cad$^-$ cells cultured in the presence or absence of NCT503 (**d**) Representative fluorescence images of intracellular ROS in E-cad$^+$ and E-cad$^-$ cells cultured in the presence or absence of 15 μM NCT503 and 1 mM NAC. (**e**) Quantified levels of ROS in E-cad$^+$ cells cultured in the presence or absence of NCT503 and NAC. Statistical analyses were conducted with unpaired two-tailed t test. Error bars indicate SEM (ns: not significant; *: $p < 0.05$; **: $p < 0.01$; ***: $p < 0.001$; ****: $p < 0.0001$).

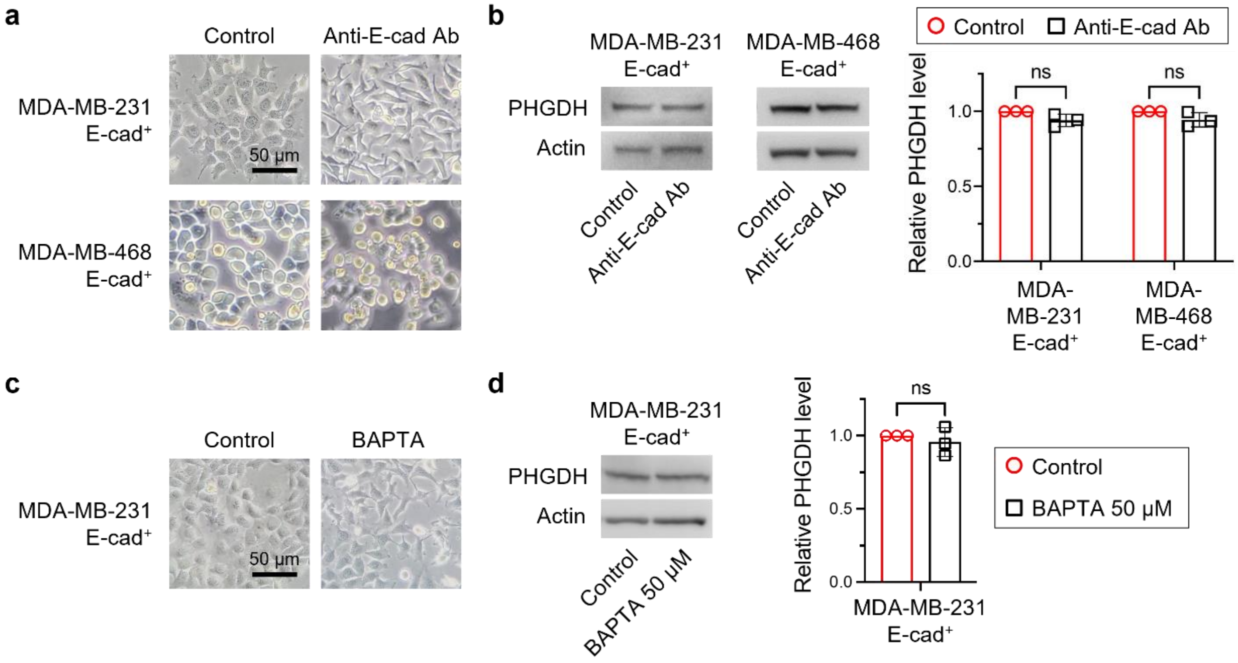


**Extended Data Fig. 4. Physical intercellular interaction is not necessary for E-cad-mediated PHGDH upregulation.**

(**a**) Representative images of E-cad$^+$ MDA-MB-231 and MDA-MB-468 cells treated with or without E-cad functional blocking antibody (HECD-1, 10 µg/ml) for 48 h. (**b**) Representative immunoblot analysis of PHGDH in E-cad$^+$ MDA-MB-231 and MDA-MB-468 cells. (**c**) Representative images of MDA-MB-231 cells treated with or without BAPTA (50 µM). (**d**) Representative immunoblot analysis of PHGDH in E-cad$^+$ MDA-MB-231 cells treated with or without BAPTA (50 µM) for 48 hr. Statistical analyses were conducted with unpaired two-tailed t test. Error bars indicate SEM (ns: not significant)

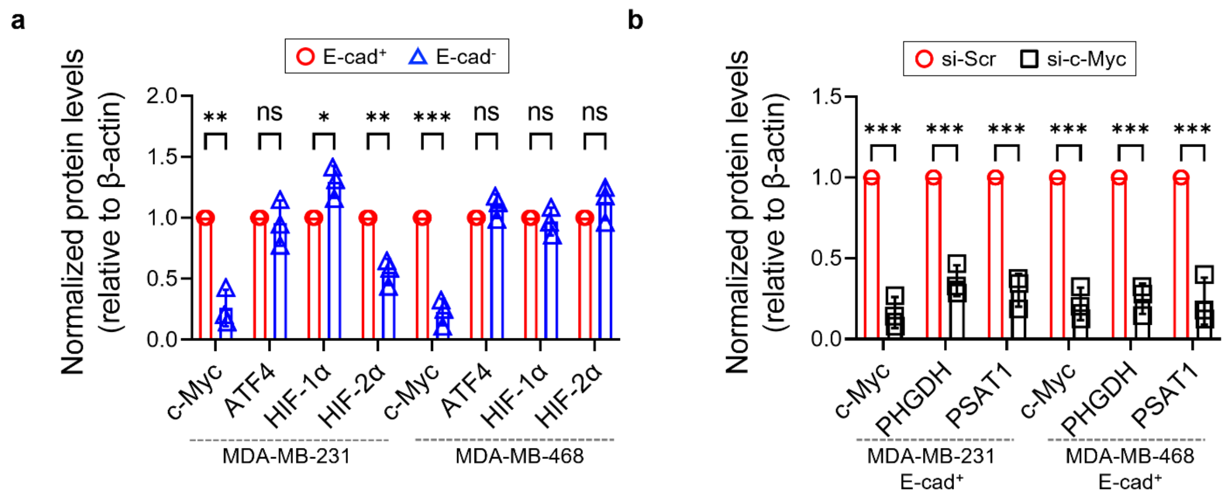


**Extended Data Fig. 5. c-Myc regulates the SSP in E-cad$^+$ breast cancer cells.**
(**a**) Normalized protein levels of E-cad$^+$ and E-cad$^-$ MDA-MB-231 and MDA-MB-468 cells. (**b**) Normalized protein levels of si-Scr and si-c-Myc E-cad$^+$ MDA-MB-231 and MDA-MB-468 cells. Statistical analyses were conducted with unpaired two-tailed t test. Error bars indicate SEM (ns: not significant; *: $p < 0.05$; **: $p < 0.01$; ***: $p < 0.001$).

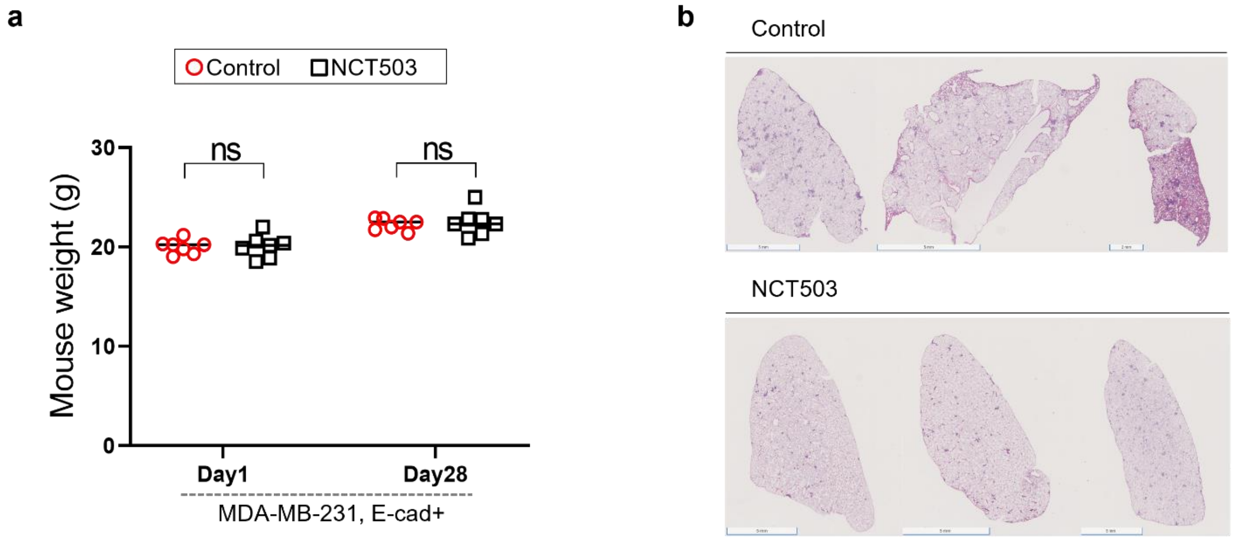


**Extended Data Fig. 6. PHGDH inhibitor treatment decreases metastasis.**
(**a**) Mice weight of first and last day of treatment. (**b**) Lung H&E staining for control (vehicle) and NCT503-treated (40 mg/kg/daily) mice. Statistical analyses were conducted with unpaired two-tailed t test. Error bars indicate SEM (ns: not significant).

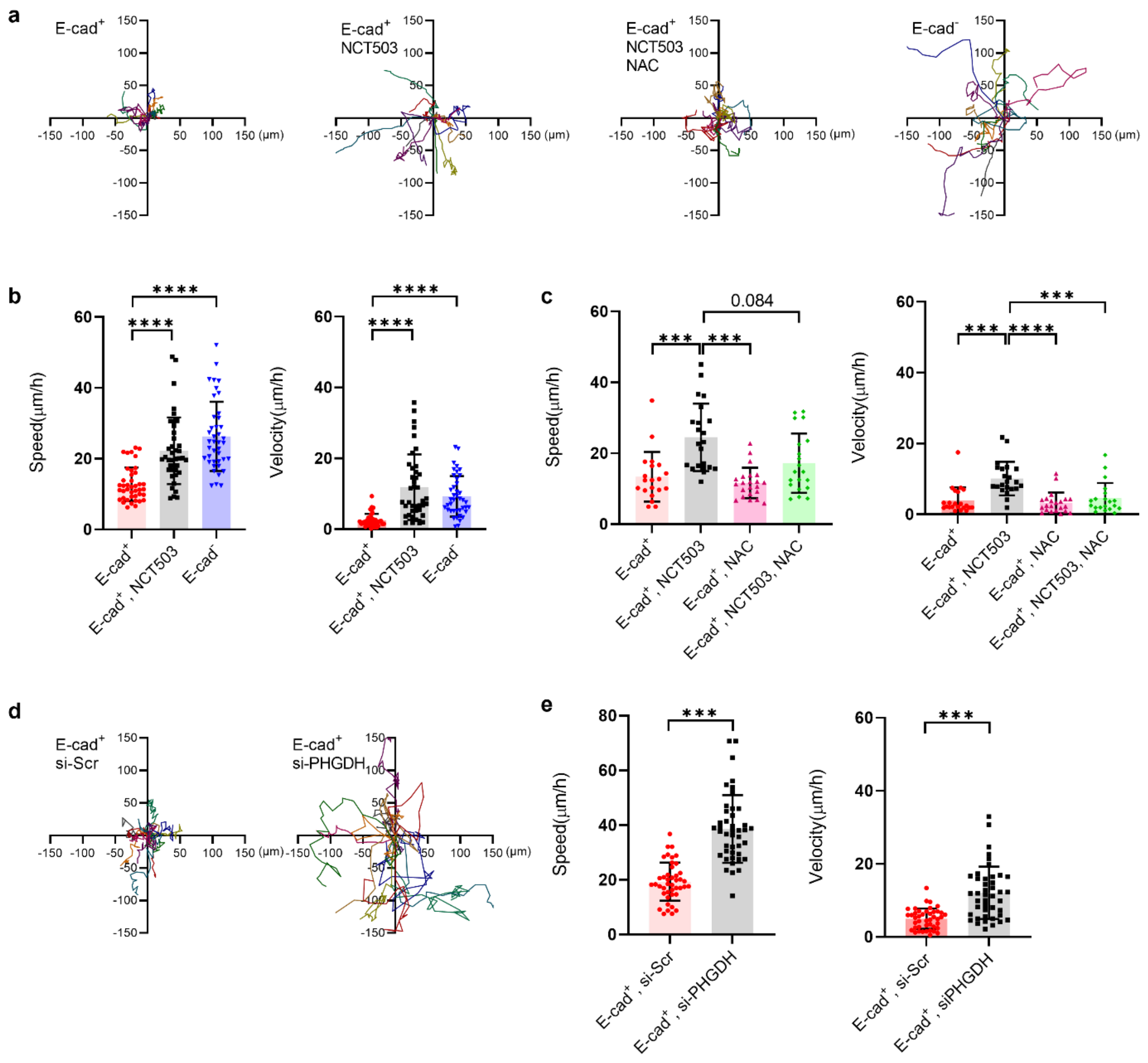


**Extended Data Fig. 7. PHGDH inhibition increases the migratory behaviors of E-cad$^+$ breast cancer cells.**

(**a**) Track overlay of 12 individual E-cad$^+$ and E-cad$^-$ MDA-MB-231 cells over 12 hr. (**b**) Analysis of 2D migratory behaviors of MDA-MB-231 cells with NCT503. (**c**) Analysis of 2D migratory behaviors of E-cad$^+$ MDA-MB-231 cells with NCT503 and/or NAC. (**d**) Track overlay of 12 individual E-cad$^+$ MDA-MB-231 cells with si-Scr or si-PHGDH over 12 hr. (**e**) Analysis of 2D migratory behaviors of E-cad$^+$ MDA-MB-231 cells with si-Scr or si-PHGDH. Statistical analyses were conducted with unpaired two-tailed t test. Error bars indicate SEM (***: $p < 0.001$; ****: $p < 0.0001$).

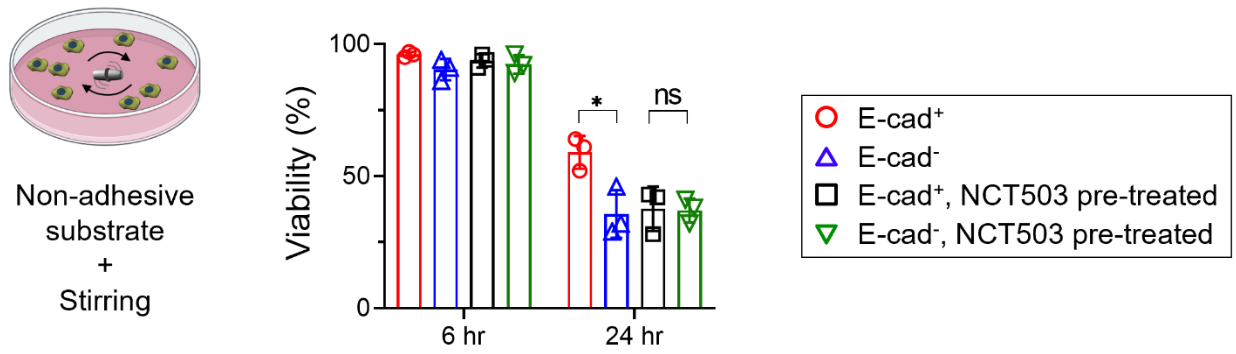


**Extended Data Fig. 8. PHGDH inhibitor treatment makes E-cad$^+$ breast cancer cells more vulnerable to cell death during circulation.**

Viability of E-cad$^+$ and E-cad$^-$ MDA-MB-231 cells after 24-hr culture in circulation. Statistical analyses were conducted with unpaired two-tailed t test. Error bars indicate SEM (ns: not significant; *: $p < 0.05$).
